# Spatial and Temporal Influences on Discrimination of Vibrotactile Stimuli on The Arm

**DOI:** 10.1101/497552

**Authors:** Valay A Shah, Maura Casadio, Robert A Scheidt, Leigh A Mrotek

## Abstract

Body-machine interfaces (BMIs) provide a non-invasive way to use and control external devices such as powered wheelchairs. Vibrotactile stimulation has been proposed as a way for BMIs to provide device performance feedback to the user, thereby reducing visual demands of closed-loop control. To advance the goal of developing a compact, multivariate vibrotactile display for BMIs, we performed two 2-alternative, forced choice experiments to determine the extent to which vibrotactile perception might vary across multiple stimulation sites. The first experiment assessed vibrotactile discrimination of sequentially presented stimuli within each of four dermatomes of the arm (C5, C7, C8, T1) and on the ulnar head. The second compared discrimination when pairs of vibrotactile stimuli were presented simultaneously vs. sequentially both within and across dermatomes. Although the first experiment found small but statistically significant differences across dermatomes C7 and T1, discrimination thresholds at the other three locations did not differ one from another or from those at either C7 or T1. These results suggest that stimuli applied to each of the sites may be able to convey approximately the same amount of information. The second experiment found that sequential delivery of vibrotactile stimuli resulted in better discrimination than simultaneous delivery, independent of whether the pairs were located within the same dermatome or across dermatomes. Taken together, our results suggest that the arm may be a viable site to transfer multivariate information via vibrotactile feedback for body-machine interfaces. However, user training may be needed to overcome the perceptual disadvantage of simultaneous vs. sequentially-presented stimuli.

## INTRODUCTION

Even the simplest of actions – such as reaching out toward a coffee mug – typically require the brain to integrate information from multiple senses to plan and execute the motor commands required to accomplish the task [c.f., (Scott, 2004)]. In healthy individuals, vision (to locate the desired object relative to the hand) and intrinsic proprioception (to sense body configuration and movement) play key roles in these processes (Sober & Sabes, 2003). Unfortunately, diseases such as Parkinson’s Disease (Vaugoyeau, Viel, Assaiante, Amblard, & Azulay, 2007), multiple sclerosis (Gandolfi, et al., 2015) and neuromotor injury [e.g. spinal cord injury (Crewe & Krause, 2009), stroke (Dukelow, et al., 2009)], can interrupt sensory feedback pathways that normally contribute to the accuracy and coordination of movements [c.f., (Sainburg, Poizner, & Ghez, 1993; Sainburg, Ghilardi, Poizner, & Ghez, 1995)]. Recent efforts in the development of body-machine-interfaces (BMIs) have sought to mitigate sensorimotor impairments caused by disease and injury by using technology to compensate for sensory and/or motor deficits (Mussa-Ivaldi & Miller, 2003). Here, we focus on the mitigation of somatosensory deficits related to the sense of body configuration and movement (i.e., proprioceptive deficits).

Various approaches to the development of sensory BMIs have included auditory, haptic, vibratory, and electro-stimulation [c.f., (Mussa-Ivaldi & Miller, 2003; Casadio, Ranganathan, & Mussa-Ivaldi, 2012)]. Vibrotactile feedback is an inexpensive and noninvasive method for displaying detailed sensory information to a user without taxing visual or auditory attention, while also improving environment exploration and motor control (Sienko, Balkwill, Oddsson, & Wall, 2008; Krueger, Giannoni, Shah, Casadio, & Scheidt, 2017; Tzorakoleftherakis, Murphey, & Scheidt, 2016; Cipriani, D’Alonzo, & Carrozza, 2012; An, Matsuoka, & Stepp, 2011). Vibrotactile perception and stimulation have been studied widely and have advanced the development of vibrotactile displays (Cholewiak, 1999; Cholewiak & Collins, 2003; Wentink, Mulder, Rietman, & Veltink, 2011; Verrillo, 1985; Harris, Arabzadeh, Fairhall, Benito, & Diamond, 2006; Tannan, Simons, Dennis, & Tommerdahl, 2007). Perception of vibrotactile stimuli depends on location of stimulation, delivery method, and cognitive ability of the user (Cholewiak, 1999; Cholewiak & Collins, 2003). Many of these prior studies focused on the hand and the digits as targets of stimulation because they are the most dexterous and are used to interact with the environment regularly (Verrillo, 1985; Tannan, Simons, Dennis, & Tommerdahl, 2007; Harris, Arabzadeh, Fairhall, Benito, & Diamond, 2006; Morley & Rowe, 1990; Post, Zompa, & Chapman, 1994). The arm may also be an ideal site to apply vibrotactile cues as the hand and digits could remain dexterous for object manipulation. However, few investigations examined perception and discrimination of vibrotactile stimuli on the arm, especially for locations other than the volar forearm. Characterizing vibrotactile discriminability of the arm will determine the acuity with which a BMI can present relevant performance information variables to the user.

The volar forearm has been the chosen stimulation site for some studies (Mahns, Perkins, Sahai, Robinson, & Rowe, 2006; Post, Zompa, & Chapman, 1994; Cholewiak & Collins, 2003; Lamore & Keemink, 1988; Morioka, Whitehouse, & Griffin, 2008), but other locations of the arm were not studied. Mahns et al. (2006) investigated vibrotactile frequency discrimination in glabrous versus hairy skin and reported discrimination thresholds of the fingertip (27.2 Hz) and the forearm (33.9 Hz), with the fingertip showing about 20% better discrimination. The density of Pacinian Corpuscles (PCs, the mechanoreceptor most sensitive to vibrations of 80-1000 Hz) is much higher in the glabrous skin of the fingertip than the hairy skin of the forearm (Hunt, 1974; Burgess, 1973), and thus, the index finger has better vibrotactile frequency discriminability than the forearm, as shown by Mahns et al (2006).

In addition to receptor density, organization of the tactile sensory area in the brain also influences our ability to discriminate tactile stimuli because information channeled from different areas might interfere with each other (Hoechstetter, et al., 2001). Non-human primate studies have shown that afferent signals from the dermatomes are projected onto somatosensory cortex in a way that preserves the arrangement of the spinal segments (Woolsey, Marshall, & Bard, 1943; Werner & Whitsel, 1968). Woolsey et al. found that dermatomes C2-C8 are projected to large and overlapping areas of the cortex, whereas dermatomes T1-T12 are mapped onto a single small location that shows minimal overlap with the projection of the cervical dermatomes. This projection pattern of dermatomes in the cortex is likely to be similar in humans as well (Penfield & Boldrey, 1937; Eickhoff, Grefkes, Zilles, & Fink, 2006) and should therefore influence our ability to discriminate vibrotactile stimuli (Duncan & Boynton, 2007). In the present study, we plan to test the discrimination thresholds in various locations in the arm. Perception and discriminability of vibrotactile stimuli dictates the acuity with which we can sense the information encoded by vibrotactile feedback. We expect that by using a similar range of vibration frequency as Mahns et al. (2006), we will find varying discrimination thresholds in the tested locations on the forearm. Characterizing discrimination thresholds within the arm will advance the development of BMIs utilizing vibrotactile feedback by allowing maximal information transfer to the body.

Discrimination of vibrotactile stimuli is influenced not only by the stimulation site but additionally by the method in which the stimuli are delivered (i.e. whether they are delivered sequentially or simultaneously, over one or more locations). Tannan et al. (2007) reported that when discriminating vibrations on the hand, people have the same discriminability regardless of whether stimuli are delivered sequentially or simultaneously. However, as the distance between the two stimuli decreased (below the two-point discrimination distance), the discriminability of simultaneous stimuli decreased, whereas for the sequential stimuli, it remained unaffected. The discriminability varies substantially for simultaneous vibrotactile stimulations based on the distance between stimulation site.

Perceptual decision making also plays a large role in the discrimination of two vibrotactile stimuli, because both memory and attention are involved in the central processes that compare two vibrotactile stimuli (Heekeren, Marrett, & Ungerleider, 2008). in the central nervous system, sensory stimuli are represented as neuronal responses, which are used to compute a decision and response on the discrimination of the stimuli. Discriminating between two sequential stimulations requires the first stimulus to be stored in working memory, which can later be accessed to compare against a second stimulus (Romo, Hernandez, Zainos, Lemus, & Brody, 2002). Additionally, stimuli stored in working memory are subject to neural noise, and the neuronal response can degrade if it is not accessed right away, leading to worse discriminability (Harris, Miniussi, Harris, & Diamond, 2002; Gallace, Tan, Haggard, & Spence, 2008). In the case of simultaneous stimuli, attentional resources are divided between the two delivery locations for successful perception of the stimuli (Connell & Lynott, 2012). Dividing attention across multiple sensory inputs weakens the integration of those inputs in the decision-making process. More so, the convergence of the sensory representation of the two stimuli is weakened before the decision-making process in the brain (Wyart, Myers, & Summerfield, 2015). Thus, we expect that the discriminability of vibration stimuli applied to the forearm will depend on the temporal profile of those stimuli.

In this study, we sought to investigate and understand the influence of spatiotemporal features of stimulus delivery on discriminability of vibrotactile stimuli. We performed a series of two-alternative forced choice experiments that examined discrimination of sequential and simultaneous vibrotactile stimuli within and across dermatomes of the arm and hand. The experiments were designed to test two hypotheses. First, based on cortical representation of dermatomes and mechanoreceptor density, we hypothesized that the acuity of vibrotactile discrimination differs across dermatomes of the arm. Second, based on influence of attention and perceptual decision making, we hypothesized that discrimination of vibrotactile stimuli is additionally influenced by stimulus delivery method (i.e., sequential vs. simultaneous). Our investigations characterized the just noticeable differences of vibrotactile stimuli delivered to the arm and identified the effects of stimulus timing on vibrotactile perception. The results will advance the development of BMIs by enhancing the utility of information encoded within vibrotactile feedback, such as force sensation for upper extremity amputees (An, Matsuoka, & Stepp, 2011), kinesthetic information for survivors of stroke (Krueger, Giannoni, Shah, Casadio, & Scheidt, 2017), or the offloading of visual attention in spinal cord injury patients (Cincotti, et al., 2007).

## MATERIAL and METHODS

### Participants

Thirty neurologically intact participants (14 females), ranging in age from 19 to 29 years (22.9 ± 2.05 yrs, mean ± SD), with no known cognitive deficits or tactile deficits of the arm were recruited from the Marquette University community. Participants gave written, informed consent to participate in one of two experiments. All experimental procedures were approved by Marquette University’s Institutional Review Board in full accordance with the Declaration of Helsinki.

### General Experimental Setup

Participants were seated with the dominant arm relaxed on a one-inch thick memory foam pad on top of a table. The elbow was oriented at 90 degrees relative to the torso. Vibrotactile stimuli were delivered to the forearm and hand via 10mm eccentric rotating mass vibration motors (“tactors”: Precision Microdrives Ltd, Model # 310-117) with an operational frequency range of approximately 60-240 Hz, which was coupled to an amplitude range of 0.5-2.4 G. For simplicity, we chose to represent vibrotactile stimuli intensity in terms of frequency even though the amplitude of the vibration covaries with frequency in the eccentric rotating mass tactors used in this study. The tactors were powered and controlled using drive circuitry that was interfaced to a portable laptop computer running a custom script within the Matlab R2017a computing environment (MathWorks Inc., Natick MA). Tactors could be placed on five locations: dermatome C5, C7, C8, T1, or the ulnar head (UH), a boney prominence within the projection of dermatome C8. Figure 1 shows the dermatomes of the arm and the approximate locations of the testing sites. Vibration motors were fixed to the arm via Transpore tape (3M Inc.).

**Fig 1.**
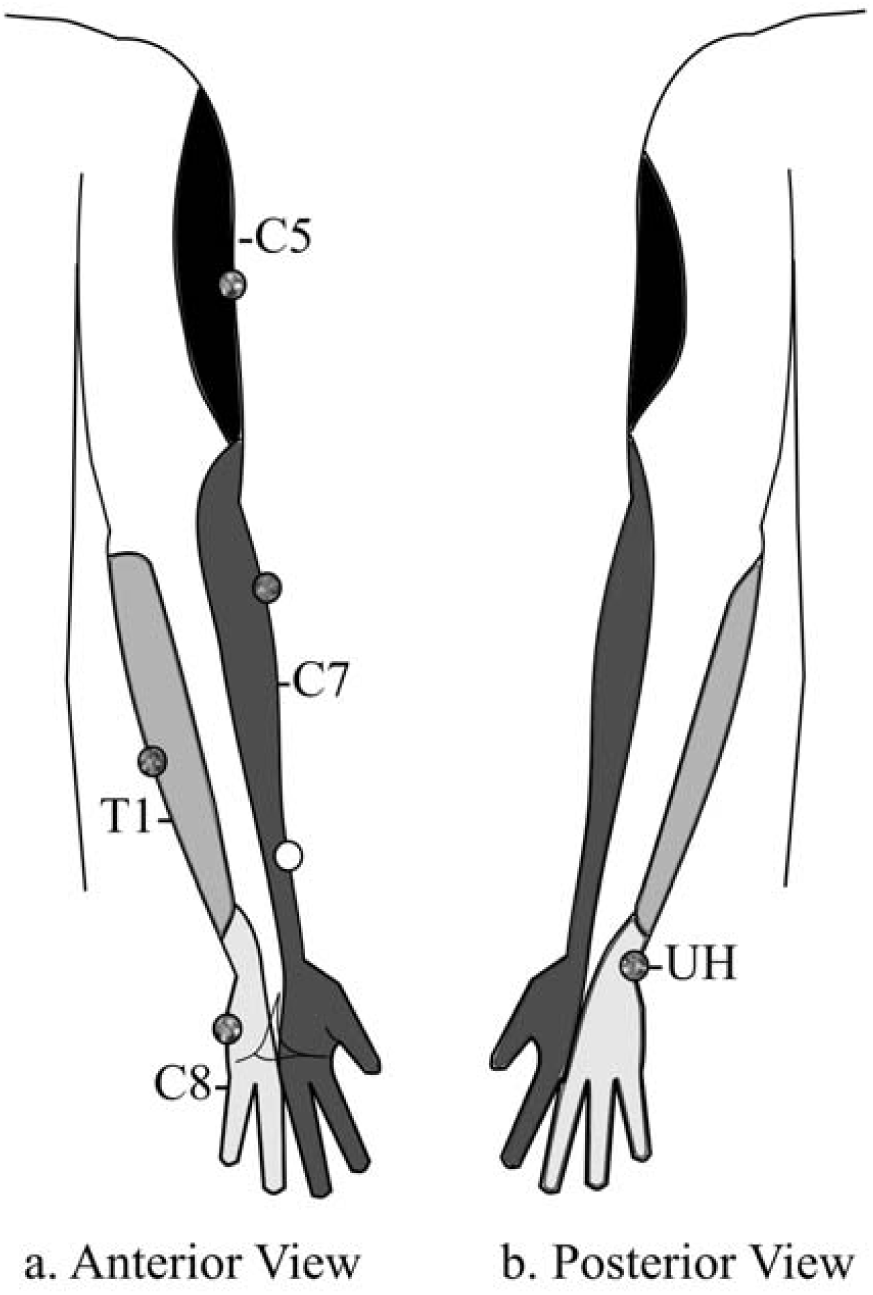
Mechanoreceptors within the arm and hand send afferent projections to one or more segments of the spinal cord through the Dorsal Root Ganglia. The dermatomes of the arm (the domains of origin of those projections) are labeled according to their target cord segment, and are marked by the *shaded regions*. The *white shaded regions* are areas of major dermatomal overlap, i.e. more than 1 spinal cord segment can innervate that region. **a)**. The anterior view of the arm, showing dermatomes, C5, C7, C8, and T1. **b)**. The posterior view of the arm, showing dermatomes and the Ulnar Head. The *gray markers* indicate the placement of the tactor motors on the arm in experimental 1 and 2. The *white marker* indicates the placement of the second tactor during the C7-C7 pair of experimental 2. Adapted from Lee et al (2008).

### Constant Stimuli Protocol

We conducted a series of two, 2-alternative, forced choice experiments (2AFC) using the method of constant stimuli (Gescheider, 1997) to determine the difference threshold of vibrotactile stimuli for each participant under various testing conditions. The 2AFC protocol presented participants with a series of 110 stimulus pairs, each comprised of a standard stimulus value that remained fixed throughout the experimental session, and a probe stimulus value that varied across stimulus pairs. The standard stimulus for our experiments was set to a frequency (186 Hz) in the middle range of the Pacinian Corpuscle’s frequency sensitivity band. The probe stimulus included five frequencies below and five above the standard stimulus value (ranging from 100-235 Hz).

For experiment 1, a single tactor was used to present two sequential vibrations at each of several different locations. Participants indicated which stimulus, first or second, they perceived to be greater in vibrotactile intensity. For experiment 2, two tactors were used to present pairs of vibrations (sequentially or simultaneously) across pairs of stimulation sites. In this case, the participant indicated the location of the stimulus perceived to be greater in vibrotactile intensity.

### Presentation of Stimuli

#### Sequential

During the sequential presentation of stimuli, the first vibrotactile stimulus was delivered for 750 ms, followed by a 750 ms pause, and then the second stimulus presentation for 750 ms (Figure 2).

**Fig 2.**
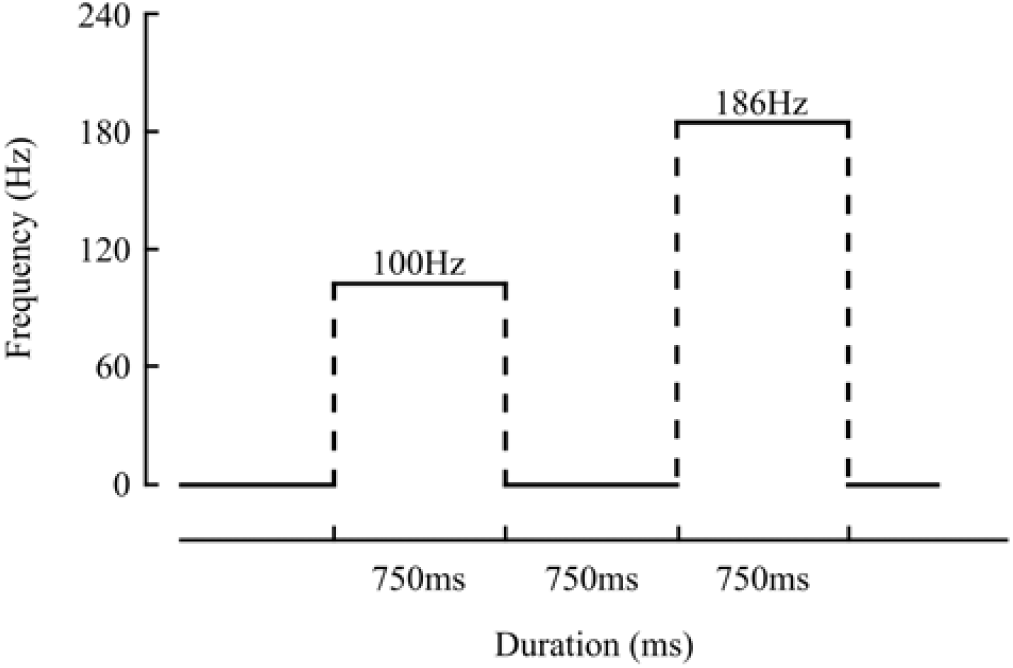
An exemplar schematic of the sequential delivery of vibrotactile stimuli. In this example, the probe stimulus is delivered first for 750 ms, then a pause for 750 ms, followed by the standard stimulus for 750 ms. If correctly discriminated, the participant would identify the second stimulus as being greater in intensity.

#### Simultaneous

During the simultaneous presentation of stimuli, both vibrotactile stimuli were provided at the same time for a duration of 750 ms. This presentation method was only used for experiment 2, wherein two tactors delivered vibrotactile stimuli to several location pairs.

### Experiment 1: Discrimination Thresholds for Sequential Stimulations in Dermatomes of the Arm and Hand

Fifteen participants (6 Females; mean age 23.7 ± 2.5 yrs) volunteered to participate in three experimental sessions, lasting approximately 60 minutes each, spaced at least 24 hours apart. Each session consisted of five blocks of 2AFC trials. During each block, one tactor was attached to the arm at one of five arm locations: C5, C7, C8, T1, or UH (Fig 1: gray markers). The vibration discrimination threshold was tested using sequential stimuli presentation. Participants completed 110 trials during each block (11 probe stimuli repeated 10 times each), wherein they verbally indicated which of the two stimuli they perceived to be more “intense”, regardless of whether they interpreted stimulus intensity to refer to stimulus magnitude or frequency. The ordering of standard and probe stimuli presentation (i.e. which stimulus was presented first) was pseudorandomized across trials. Testing locations were also pseudorandomized across participants and sessions to minimize potential order effects.

### Experiment 2: Sequential versus Simultaneous Stimulation Within and Across Dermatome Pairs

Fifteen participants (8 Females; mean age 22.2 ± 1.2 yrs) volunteered to participate in a single experimental session, lasting approximately 90 minutes. The session consisted of eight blocks of 2AFC trials. During each block, one of four dermatomal pairs were tested using either sequential or simultaneous stimuli presentation: within a dermatome (C7-C7), and across dermatomes (C7-C5, C7-UH, and C7-T1). One tactor was always placed on dermatome C7 at the location marked by the gray C7 marker in Figure 1. A second tactor was attached to the other indicated location. The two tactors were always placed at least 8cm apart, as results of pilot testing on the propagation of vibrations across the arm revealed that cross-coupling of vibration across stimulation sites was negligible with tactor separations greater than 6 cm [data not shown; see also (Krueger, Giannoni, Shah, Casadio, & Scheidt, 2017; Cipriani, D’Alonzo, & Carrozza, 2012)]. Participants completed 110 trials during each block, where they verbally indicated which of the two tested locations received the more “intense” stimulation. The ordering of standard and probe stimuli (i.e. which stimulus was presented at which location) was pseudorandomized across trials. Block presentation order [i.e., the eight combinations of stimulation delivery method (sequential / simultaneous) and sites (dermatomal pairs)] were also pseudorandomized across participants and blocks to minimize potential order effects.

### Data Analysis

Verbal responses were converted into probabilities of indicating each probe stimulus as being greater in intensity than the standard stimulus. For each participant and each testing block, psychometric functions were fit to the probability data as a function of probe stimulus frequency using the cumulative normal distribution (Eq 1).

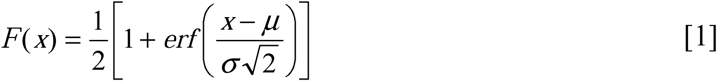

where, F(x) is the predicted probability, x is the probe frequency, μ is the mean of the underlying decision process modelled as a normal distribution, σ is the standard deviation of that normal distribution, and the *erf* is the cumulative normal function. Curve fitting was performed using the MatLab function (`fminsearch`) to find the μ and σ values that minimized the sum of squared error between the predicted and actual response probabilities. The discrimination threshold was defined as one standard deviation of the underlying normal distribution (i.e. the σ found by `fminsearch`). As we found no significant effect of sessions for experiment 1 (see Results), discrimination thresholds were averaged across the three sessions for each tested location, to provide one discrimination threshold per participant. For both experiments 1 and 2, we report the mean discrimination threshold averaged across participants within blocks.

### Statistical Hypothesis Testing

Motivated by the observation that the density of cutaneous mechanoreceptor varies across the body (Hunt, 1974), we first sought to test the extent to which discrimination thresholds for vibrotactile stimuli might vary across locations of the arm and hand (Experiment 1). Specifically, we used two-way ANOVA and post-hoc, Bonferroni-corrected, paired samples t-test to verify the hypothesis that mean vibrotactile discrimination thresholds vary across locations on the arm and hand.

Motivated by the consideration that discrimination of sequential vibrotactile stimuli involves aspects of working memory not required for simultaneously presented stimuli, we sought to test the hypothesis that discrimination thresholds would vary between sequential and simultaneously presented stimuli, both within and across dermatomes (Experiment 2). We used two-way ANOVA and post-hoc, Bonferroni-corrected, paired samples t-test to compare discrimination threshold (the dependent variable) across delivery methods (sequential or simultaneous) and across location of stimulus delivery (within or across dermatomes).

All analyses were performed with SPSS Statistics 24 (IBM Corp, Armonk, NY). Statistical significance was set at the family-wise error rate of α = 0.05.

## RESULTS

This study used eccentric rotating mass tactors to examine the psychophysics of vibrotactile perception within and across dermatomes of the arm and hand in 30 neurologically healthy participants. All participants were attentive throughout their experimental session, and all responded to stimuli in a timely fashion. Each session lasted about 60 minutes in Experiment 1 and about 90 minutes in Experiment 2, depending on participant response times (each trial lasted about 3-5 s).

### Experiment 1: Discrimination Thresholds for Sequential Stimuli Applied at Single Locations in Dermatomes of the Arm and Hand

In the first set of experiments, we tested the extent to which vibrotactile discrimination thresholds vary across dermatomes of the arm and hand. Figure 3 (*left*) depicts response probabilities calculated from a single block of discrimination trials from dermatome C7, performed by one participant. When the probe stimulus frequency was markedly lower than that of the standard stimulus, the participant reliably identified the standard as more intense than the probe [i.e., P (probe > standard) was close to 0]. By contrast, when the probe stimulus frequency was markedly higher than that of the standard, the participant was much more likely to identify the probe stimulus as more intense. When the probe stimulus frequency was close to that of the standard stimulus, the participant was less reliable in correctly identifying which stimulus was more intense. We fit the cumulative normal function (Eq 1) to the observed likelihood data in order to obtain estimates of µ and σ from the underlying normal model of the perceptual decision process. Figure 3 (*right*) presents the psychometric curves obtained from all five testing locations from the same participant. Dermatome C5 is traced by the blue curve (174.27 ± 35.87 Hz; µ ± σ of the underlying normal distribution), dermatome C7 by the red curve (186.38 ± 19.01 Hz), dermatome C8 by the orange curve (193.09 ± 46.69 Hz), dermatome T1 by the green curve (189.29 ± 64.42 Hz), and the ulnar head by the purple curve (181.16 ± 34.95 Hz). Here, the curve for dermatome C7 had the steepest slope (smallest σ) whereas dermatome T1 had the shallowest slope (greatest σ). Thus, this participant was better at discriminating between vibrotactile stimuli presented sequentially on dermatome C7 than the same stimuli presented on dermatome T1. Discrimination thresholds for sequential stimuli applied to dermatomes C5, C8, and the ulnar head fell between the bounds established by dermatomes C7 and T1.

**Fig 3.**
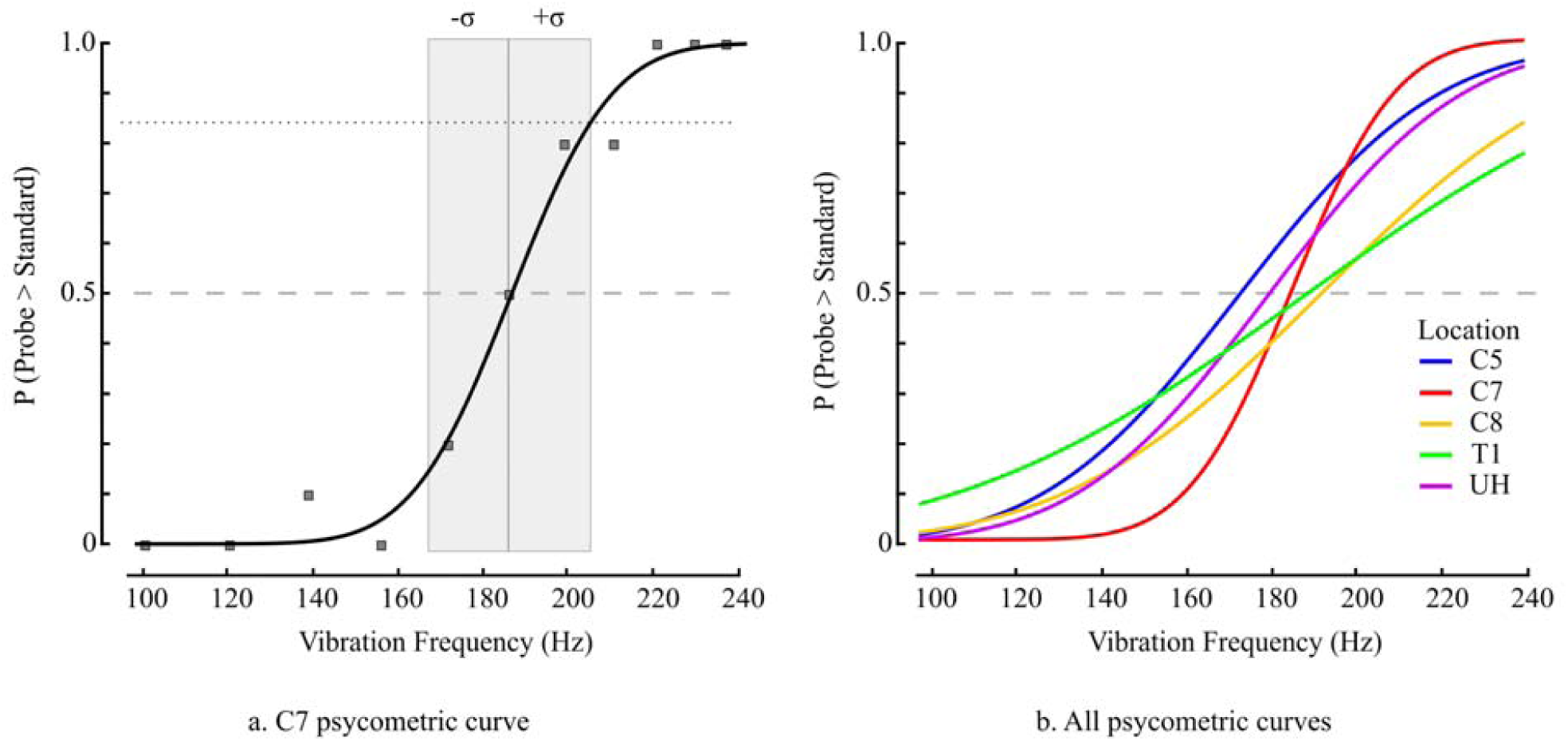
**a)**. Assessment of vibrotactile perception at dermatome C7 for a participant. *Gray Squares* indicate the observed fraction of trials at each probe frequency where the participant indicated that she perceived the probe stimulus as more intense than the standard stimulus. *Black sigmoid curve*: the psychometric (cumulative normal) function that was fit to the observed probability data. *Gray Shaded Region:* the discrimination threshold defined as one estimated standard deviation (here, ±19.01 Hz) from the estimated mean (186.38 Hz) of the underlying normal distribution. The upper bound of the box crosses the sigmoid at approximately P = 0.84 (*Gray dotted line)*. *Gray Dashed Line:* the point of subjective equality (i.e., P = 0.5). **b)**. Best-fit cumulative normal functions for the five testing locations for the same participant. Dermatome D7 has the best discrimination threshold, while dermatome T1 has the worst.

The results presented in Figure 3 were representative of the study population as a whole (Fig 4). Two-way ANOVA found that vibrotactile discrimination differed significantly across stimulation sites (F_4,56_ = 6.801, p = 0.0002), but not across session (F_2,28_ = 1.212, p = 0.313). Post-hoc testing revealed that this effect was due to better vibrotactile discrimination on dermatome C7 [32.78 ± 4.73 Hz (mean ± SEM)] vs. dermatome T1 (43.25 ± 5.48 Hz, t_14_ = 5.22, p = 0.0001). Vibrotactile discrimination thresholds on dermatomes C5 (36.88 ± 4.23 Hz), C8 (37.96 ± 4.58 Hz), and the Ulnar Head (34.70 ± 4.03 Hz) did not differ significantly from each other or from those on dermatomes C7 or T1 (p > 0.05 in all cases). Across participants, the average difference in discrimination thresholds between dermatomes C7 and T1 was 10.47 ± 1.48 Hz.

**Fig 4.**
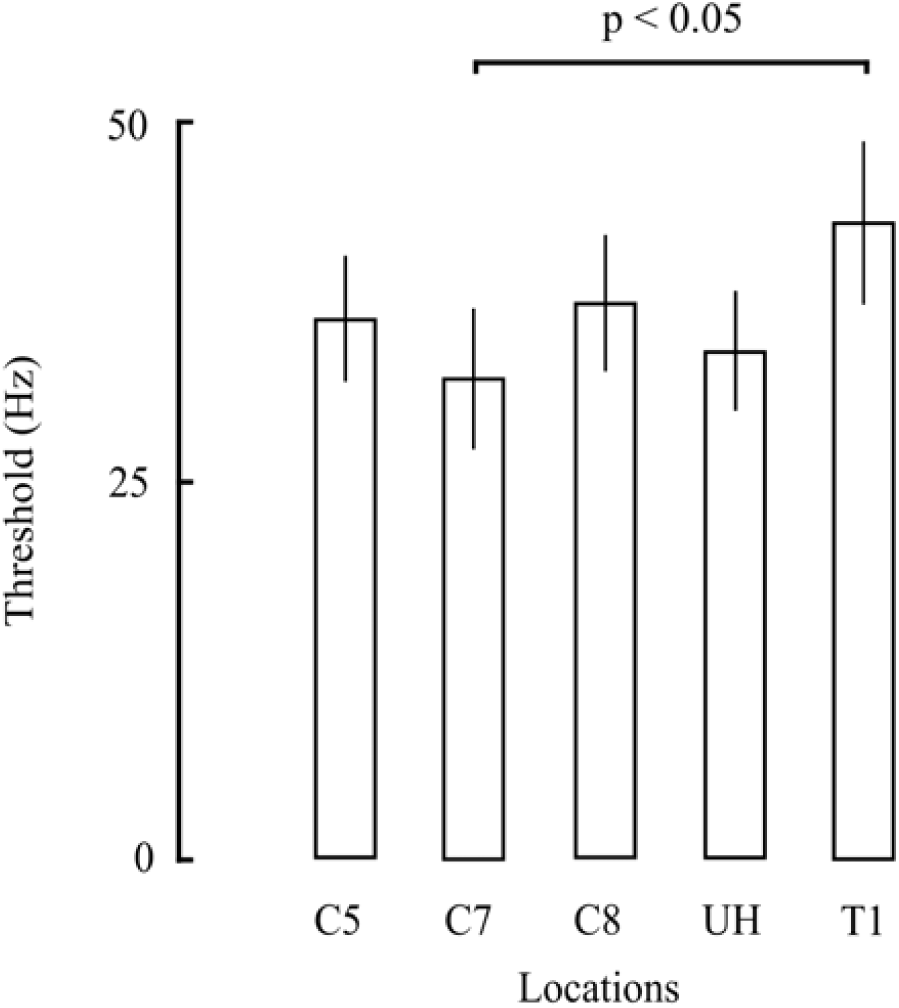
Group results from Experiment 1. Mean (± 1 SEM) discrimination thresholds across the participant population were calculated for sequential vibrotactile stimuli presented within each of the five tested locations. Dermatome C7 is significantly better at discriminating vibrotactile stimuli than dermatome T1.

**Fig 5.**
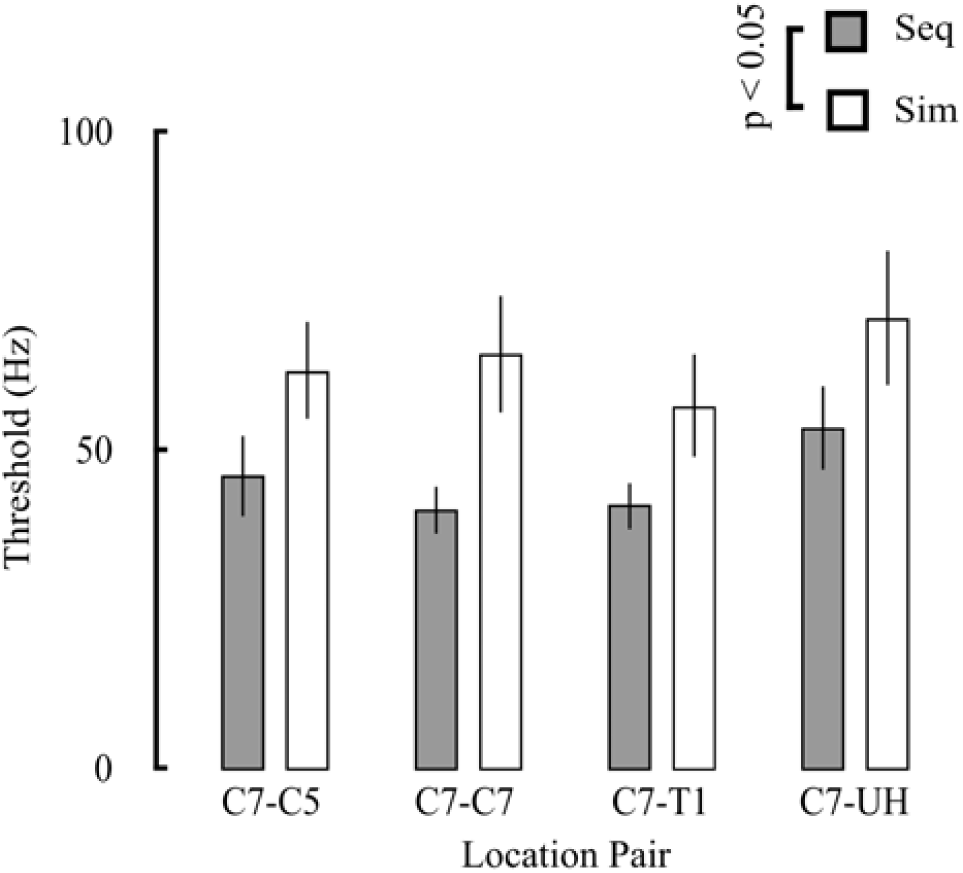
Group results from Experiment 2. Mean (± 1 SEM) discrimination thresholds were calculated for sequentially (gray bars) and simultaneously delivered (white bars) vibrotactile stimuli at stimulus location pair. Sequential vibrotactile stimuli allowed for better discriminability than simultaneous stimuli.

### Experiment 2: Sequential versus Simultaneous Stimulation Within and Across Dermatome Pairs

In the second set of experiments, we examined two factors having potential to impact how the brain processes vibrotactile information in support of perceptual decision making: concurrency of stimuli (i.e., whether working memory is required to support the decision) and somatotopy of stimulus delivery (i.e., whether the two stimuli are provided within the same dermatome or across different dermatomes). Participants performed 8 blocks of 2AFC trials wherein they discriminated between two vibrotactile stimuli delivered either sequentially or simultaneously at each of four location pairs on the arm or hand; each permutation of this 2×4 experimental design was tested in separate blocks. As per Experiment 1, we fit Eq 1 to the observed response likelihood data from each block to obtain separate estimates of the mean (µ) and standard deviation (σ) of the normal model of the perceptual decision process underlying each testing condition. Two-way ANOVA found that vibrotactile discrimination thresholds varied systematically by delivery method (F_1, 113_ = 13.01, p = 0.0004), but did not depend significantly across paired stimulation sites (F_3, 113_ = 1.124, p = 0.343). Participants demonstrated better discriminability of vibrotactile stimuli with sequential delivery [45.57 ± 3.92 Hz (mean ± SEM)] than with simultaneous delivery (64.14 ± 6.54 Hz). Across participants, the difference in discrimination thresholds between delivery methods averaged 18.57 ± 7.83 Hz. The main effect found in experiment 1 did not differ significantly from the main effect found in experiment 2 (2-sample t test, t_28_ = 1.0167, p = 0.318).

## DISCUSSION

This study investigated the discriminability of vibrotactile stimuli applied either sequentially or simultaneously to various dermatomes on the arm and hand (C5, C7, C8, T1, and the UH). Based on reports of differing densities of mechanoreceptors in the hand and varying dermatomal projections in the Somatosensory Cortex (SI and SII), we hypothesized that the discrimination threshold for vibrotactile stimuli would vary across dermatomes. In support of this hypothesis, we observed a small but significant difference in vibrotactile discrimination between dermatome C7 and T1, whereas discrimination thresholds in dermatomes C5, C8, and the UH did not differ from C7, T1, or one another. The current study also tested the hypothesis that the discriminability of vibrotactile stimuli depends on whether the stimuli are delivered sequentially or simultaneously. Our results showed that the discriminability of sequentially delivered stimuli was better than that of simultaneously delivered stimuli, regardless of whether pairs of stimuli were delivered within or across dermatomes.

These results suggest that all tested stimulation sites are valid locations for vibrotactile feedback used in body-machine-interfaces (BMIs), although the impact that differing acuity in dermatomes C7 and T1 may have on information transmission through a vibrotactile display should be examined further. In addition, if vibrotactile stimuli are provided at two separate locations, sequential delivery would permit better discriminability and would allow for more information to be transferred through vibrotactile feedback while using BMIs.

### Discrimination across dermatomes – possible mechanisms

It is possible that the difference in discrimination thresholds between dermatome C7 and T1 may be attributed to differences in the cortical magnification of dermatomal projections onto the somatosensory cortex. In non-human primates, the cortical representation of dermatome C7 has a much larger area than dermatome T1 (Woolsey, Marshall, & Bard, 1943). These dermatomal representations in the somatosensory cortex likely carry over to the human brain (Penfield & Boldrey, 1937; Eickhoff, Grefkes, Zilles, & Fink, 2006), suggesting a possible mechanism for the differences we found in discrimination levels for dermatome C7 and T1 in experiment 1. Duncan and Boynton (2007) showed that, in humans, cortical magnification (i.e. the number of neurons responsible for sensing a stimulus) of the index finger is much larger than the little finger. This larger magnification causes greater tactile acuity in the index finger compared to the little finger. In our study, discrimination thresholds in the cervical dermatomes were indistinguishable, whereas dermatomes C7 and T1 differed significantly in a way that could reflect greater cortical magnification of the cervical dermatomes. Future neuroimaging work is needed to test whether cortical magnification can explain the differences in discrimination observed in this study.

A second possibility relates to potential differences in mechanoreceptor density cross the arm. Pacinian Corpuscles (PCs) are much sparser and their location is also much deeper in the epidermis in hairy skin relative to glabrous skin (Burgess, 1973). Johansson & Vallbo (1979) showed that the density of PCs is higher towards the lateral side (index finger and thumb) of the hand compared to the medial side (little finger). It is possible that this lateral to medial difference in mechanoreceptor density may also hold true for the forearm, especially since desensitization of dermatome T1 (medial arm) may occur due to frequent interactions with objects in the environment (e.g. resting the arm on a chair or a table). To our knowledge, no histological studies to date have compared mechanoreceptor density across the dermatomes of the arm.

### Influence of working memory and attention

Romo et al. (2002) reported that working memory is involved when discriminating two sequentially presented vibrotactile stimuli. In this study, responses were recorded from the Secondary Somatosensory cortex (SII) of monkeys and showed a history-based activation of neurons during a perceptual decision task, correlated to the decision of stimuli discrimination. During the delay period between the two stimuli, the neuronal response of the first stimuli was retained at the same level as the actual stimulus in the Prefrontal Cortex (PFC), but not in SII (Romo, Brody, Hernandez, & Lemus, 1999). A trace of the first stimulus was recalled into SII when the second stimulus was presented, allowing for a comparison between the intensity of the two stimuli. In the cohort of healthy participants we tested, memory encoding and recall was not responsible for any degradation in vibrotactile acuity because the results of Experiment 2 showed that sequential stimulations were discriminated with *greater* accuracy than simultaneous stimulations. If the mechanism described by Romo et al. also holds true for vibrotactile discrimination in humans, working memory systems – possibly located within PFC – facilitate levels of vibrotactile acuity that significantly exceed those observed during the presentation of simultaneous stimuli. What can explain this outcome?

Connell and Lynott (2012) report that when comparing simultaneous stimuli, attentional capacity is reduced. This follows the capacity sharing model proposed by Pashler (1994): when attentional capacity is shared or divided across stimuli, less capacity is available to be allocated to each stimulus. Attentional focus driven towards one stimulus causes interference in the perception of another stimulus (Connell & Lynott, 2012; Kaschak, Zwaan, Aveyard, & Yaxley, 2006) as attentional capacity is bounded. This interference is also linked to increased sensory noise (uninformative information) being added to stimuli (Spence, Nicholls, & Driver, 2001). When attention towards a stimulus is decreased, sensory noise increases, thereby decreasing the accurate perception of the stimulus and reducing overall ability to discriminate between two simultaneous stimuli (Mozolic, Long, Morgan, Rawley-Payne, & Laurienti, 2011).

We propose that the ability of working memory to store previous stimuli for comparison and the ability of attention to attenuate sensory noise may explain the results we obtained in the current study, where we found that sequential stimuli were better discriminated than simultaneously presented stimuli. Future studies could manipulate the duration of the interstimulus interval to quantify the effect of temporal drift or degradation in working memory (Harris, Miniussi, Harris, & Diamond, 2002; Berglund, Berglund, & Ekman, 1967; Gallace, Tan, Haggard, & Spence, 2008), which for experiment 2 was maintained below 1 second, so that little to no degradation should have occurred (Romo, Hernandez, Zainos, Lemus, & Brody, 2002).

### Implications for vibrotactile sensory augmentation

By developing an understanding of vibrotactile perception, vibrotactile feedback can be used more effectively in applications such as sensory augmentation or substitution (Bach-y-Rita, 1967). The use of vibrotactile feedback in sensory substitution has been investigated since the 1960s. Sensory substitution is a technique where deficits in one sensory modality are mitigated through the application of stimuli to another sensory modality. Previous studies have utilized the tactile sense to substitute for other impaired senses. For example, Sienko et al. (2008) provided feedback of trunk sway to users with vestibular loss, successfully reducing body sway using the vibrotactile feedback. Witteveen et al. (2015) increased accuracy of grip force and hand aperture in prosthetic users by providing information about grip force and hand aperture with vibrotactile feedback. In our earlier work (Krueger, Giannoni, Shah, Casadio, & Scheidt, 2017), we investigated the use of vibrotactile sensory augmentation for upper extremity motor control. We applied vibrotactile feedback to the non-moving arm, conveying either limb-state or error information about the moving arm, leading to significant improvements in reaching and stabilization. One reason for choosing the arm as the location for vibrotactile feedback was that the hand and digits would not be obstructed by the tactors, thereby allowing the user to manipulate objects with the contralateral arm (e.g using the contralateral arm to hold a bottle while the ipsilateral arm is opening the bottle).

The current study has implications for the location of vibrotactile feedback and the method of stimuli delivery to allow for successful transfer of information via vibrotactile feedback. For the location of the vibrotactile feedback, previous studies have selected feedback sites that are relevant to the sensory substitution application such as the around the truck when trying to reduce trunk sway or on the arm to improve motor control of the same or contralateral arm. Our current study advances the development of such sensory substitution applications that are focused on utilizing the arm as the feedback delivery site by providing a better understanding of vibrotactile perception on various locations of the arm. The results of experiment 1 characterized the levels of stimuli differences needed to accurately distinguish between two stimuli. These results can directly affect the level information that can be encoded in BMIs. We observed small but statistically significant differences across two of the observe sites, suggesting that dermatome C7 may be able to convey somewhat more information than dermatome T1. The results of experiment 2 provided insight into the method of vibrotactile stimulation, finding that sequential delivery outperforms simultaneous due to involvement of working memory and attention. Future applications of vibrotactile sensory substitution on the arm may consider using dermatomes C5, C7, or C8 (UH) as stimulation sites as they have indistinguishable discrimination thresholds, while potentially avoiding dermatome T1. Vibrotactile acuity within tactor pairs did not appear to depend on whether the pairs were located within the same dermatome or across dermatomes. Future applications may also consider limiting the extent to which attention must be divided across multiple simultaneous stimuli. Finally, it would be of interest for future studies to verify the levels of cortical magnification of dermatomes C5, C7, C8, and T1 in humans, providing insights into cortical representation of dermatomes and confirming the work in animal models by Woosley et al. (1943) and Eickhoff et al. (2006).

### Limitations

There are several potential limitations of the present study. One potential limitation is that the difference in discrimination thresholds between dermatomes C7 and T1 (Experiment 1) could affect the discrimination threshold of paired stimulations in the C7-T1 testing condition (Experiment 2). We mitigated this concern by counter-balancing the presentation of standard and probe stimuli across the two locations through pseudo-randomization. Another limitation arises from our choice to use inexpensive ERM tactors rather than more expensive devices that can decouple the frequency of vibration from its magnitude. It is unlikely however that the factors contributing to the spatiotemporal variations in vibrotactile acuity described in this study would be the result of variations in sensitivity to just one of these parameters but not the other, and so we would not expect the overall pattern of results we describe to depend on the choice of tactor technology [c.f. (Hwang, Seo, Kim, & Choi, 2013)]. Other limitations arise from our choices to include only healthy, young participants in this study, to test using only a single standard stimulus, and to test using only a single stimulus duration. Aging has shown to be a factor in perception of vibrotactile stimulations (Lin, et al., 2015) and so discrimination thresholds might vary if we conduct the same experiments in an older population. Because perception of vibrotactile stimuli appears to adhere to Weber’s law (Francisco, Tannan, Zhang, Holden, & Tommerdahl, 2008), we would expect the magnitude of discrimination thresholds to vary as a function of standard stimulus frequency. Finally, because vibrotactile perception also appears to depend on stimulus presentation time for short stimuli less than 1 second in duration (Berglund, Berglund, & Ekman, 1967), we would also expect the magnitude of discrimination thresholds to vary as a function of stimulus duration. In all of these cases, however, we would not expect the observed variations in perception across dermatomes and across temporal patterns of stimulation to change as a result of arbitrary choices in standard stimulus frequency, stimulus duration, and participant population. Future experiments of vibrotactile perception could be performed to verify these assumptions.

## ACKNOWLEDGMENTS

This work was supported by National Institutes of Health under award numbers: R01HD053727 and R15HD093086, National Science Foundation under an Individual Research and Development Plan, Marquette University Research Leaders Fellowship, a Whitaker International Program Grant, Ministry of Science and Technology, Israel (Joint Israel-Italy lab in Biorobotics “Artificial Somatosensation for Humans and Humanoids”), and EU commission FP7 People: Marie-Curie Actions (334201).

